# Early patterning followed by tissue growth establishes proximo-distal identity in *Drosophila* Malpighian tubules

**DOI:** 10.1101/2022.05.05.490769

**Authors:** Robin Beaven, Barry Denholm

**Affiliations:** Deanery of Biomedical Sciences, The University of Edinburgh, United Kingdom

**Keywords:** proximo-distal patterning, Wnt/Wingless, epithelial tubule, Dachshund, Homothorax

## Abstract

Specification and elaboration of proximo-distal (P-D) axes for structures or tissues within a body occurs secondarily from that of the main axes of the body. Our understanding of the mechanism(s) that pattern P-D axes is limited to a few examples such as vertebrate and invertebrate limbs. *Drosophila* Malpighian/renal tubules (MipTs) are simple epithelial tubules, with a defined P-D axis. How this axis is patterned is not known, and provides an ideal context to understand patterning mechanisms of a secondary axis. Furthermore, epithelial tubules are widespread, and their patterning is not well understood. Here, we describe the mechanism that establishes distal tubule and show this is a radically different mechanism to that patterning the proximal MpT. The distal domain is patterned in two steps: distal identity is specified in a small group of cells at the earliest stage of MpT development through Wingless/Wnt signalling. Subsequently, this population is expanded by proliferation to generate the distal MpT domain. This mechanism enables distal identity to be established in the tubule in a domain of cells much wider than the effective range of Wingless.

**Summary statement:** How does proximo-distal patterning occur in the epithelial Malpighian/renal tubules? Cells are patterned early by a mechanism involving Wingless/Wnt, and expand by cell proliferation to generate a distal domain.

## Introduction

Some tubular epithelia, such as nephrons of the kidney, and the oviduct, possess functional polarities along their length, in which distinct domains of cells execute specialised physiological activities. The order of these domains can be important for the emergent properties of the system, for example the sequential processing of primary urine by functionally distinct domains along the length of the nephron. For this reason, the correct and precise establishment of cells with specialised physiological activities along the length of the tube is an important aspect for how the system will ultimately function. Developmental mechanisms that establish and pattern these axes act secondary to those that the establish and pattern the main body axes of the animal. In general these mechanisms are not well described.

Insect Malpighian, or renal tubules (MpTs) are tubular epithelia with a distinct functional axis along their proximo-distal (P-D) length including distinct regions for calcium homeostasis, fluid secretion and reabsorption (Dow et al., 1994; O’Donnell and Maddrell, 1995; Sozen et al., 1997; Wessing and Eichelberg, 1978). During embryogenesis, clusters of MpT primordial cells bud out from the wall of the gut, before extending to form each of the MpTs (Skaer, 1993). Functional polarity along the P-D axis is established during this period. It occurs several hours after the patterning of anterior-posterior and dorso-ventral axes of embryo and therefore presents an interesting system to explore how such secondary axes are established and patterned during development. This provides an alternative and therefore informative comparison, for example with P-D patterning of the insect appendages which are a well-studied example of secondary axis formation. The MpTs also provide an example of how developmental patterning occurs in the context of a tubular epithelium.

In our previous paper we discovered a mechanism by which the proximal region of the MpT P-D axis is patterned in *Drosophila*. We determined that the signalling ligand Wingless/Wnt (Wg), produced in gut cells abutting the end of the proximal tubule, patterns the proximal part of the tubule. The ability of the Wg ligand to disperse from its source is essential for this function (Beaven and Denholm, 2018).

The mechanism(s) that establish distal identity is not known, and we therefore turned our attention to this problem here. We considered two putative mechanisms by which patterning of the distal segments of the MpTs could be achieved (Fig. 1). In the first, an asymmetric signal from one or other end of the MpT, would play an instructive role in defining distal tubule identity. The signal could either be a distal identity-inducing signal emanating from the distal end of the developing MpT (Fig. 1A’), or a distal identity-repressing signal emanating from the gut at the proximal MpT end (Fig. 1A’’). We have previously shown that cells derived from the tip cell lineage are a signalling hub that establish a gradient of EGF signalling activity in the distal tubule (Saxena et al., 2014), and thus might be an inductive signal for distal identity. As mentioned above, Wg emanating from the gut at the proximal tubule end induces proximal identity by regulating the expression of genes required for the differentiation of proximal cells (Beaven and Denholm, 2018). It is conceivable that Wg also acts to restrict distal identity by inhibiting the expression of genes required for the differentiation of distal cells.

**Figure 1.**
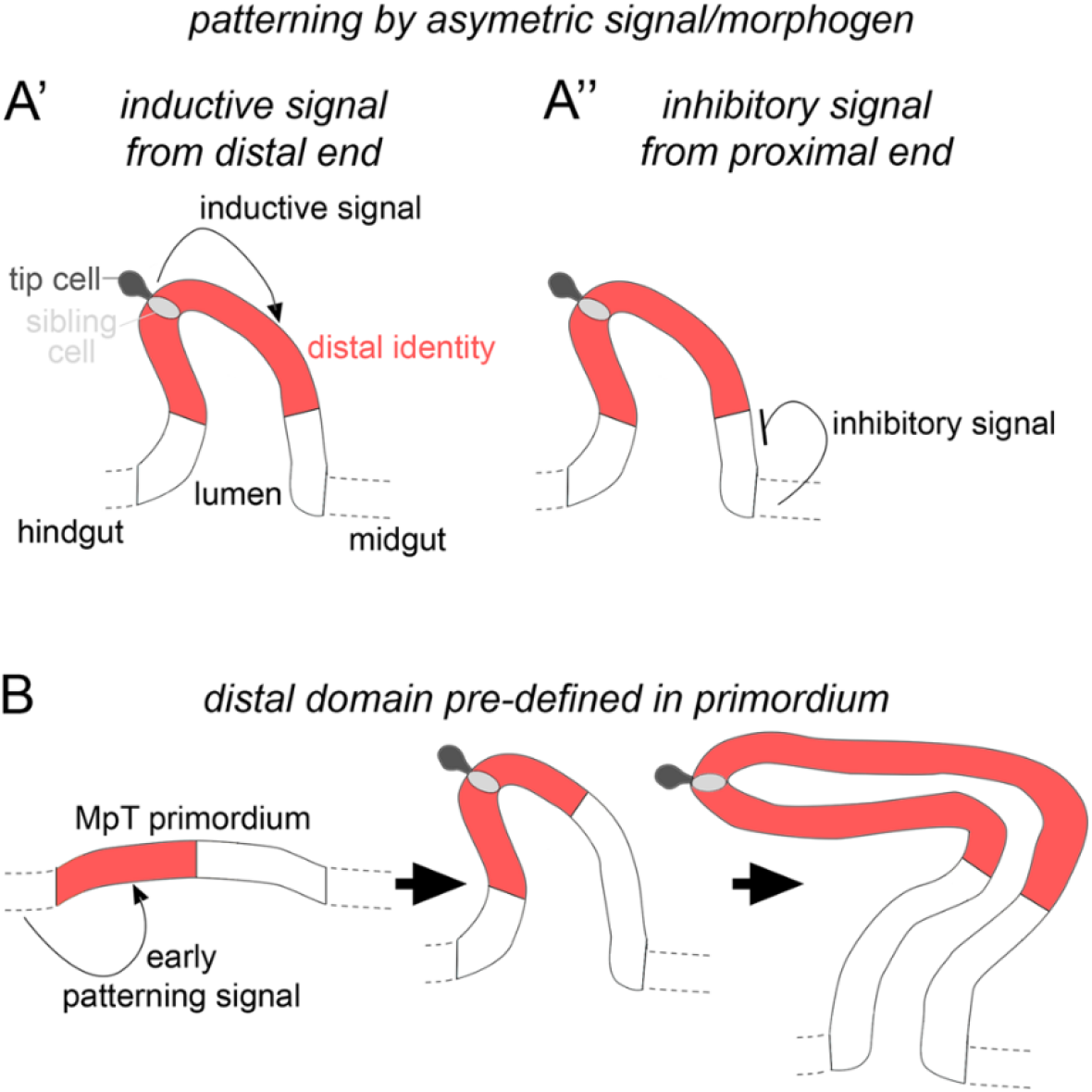
Hypothetical mechanisms for distal patterning in the developing Malpighian tubules. **(A’)** The distal tubule domain is specified by an inductive signal emanating from the distal tubule end, or **(A’’)** an inhibitory signal from the proximal tubule end. **(B)** The distal tubule domain is pre-patterned in the primordium.

The second mechanism we considered is that MpT primordia are already pre-patterned at a stage before they bud from the gut wall and that these domain identities are retained as the MpTs bud out, proliferate and elongate as a tubular structure (Fig. 1B). This would be conceptually similar to the way in which the *Drosophila* legs are patterned, with positional information defined in the leg disc which endures as the disc telescopes out to form the mature leg (Basler and Struhl, 1994; Diaz-Benjumea et al., 1994; Giorgianni and Mann, 2011; Lecuit and Cohen, 1997; Meinhardt, 1983b; refs. within Ruiz-Losada et al., 2018).

Previous studies suggested this as a plausible mechanism, and indicated that Wg could also be involved in this earlier process. There is a proneural cluster of cells within the MpT primordium, one of which gives rise to the tip-cell and sibling-cell (Hoch et al., 1994; Wan et al., 2000). Wg is expressed in the hindgut, as well as the MpT primordium from the start of tubule development (Skaer and Martinez Arias, 1992), and is required for expression of Achaete in the proneural cluster. Wg is also required for expression of Seven up and PointedP2 (Sudarsan et al., 2002; Wan et al., 2000), two distal transcription factors considered to play roles in promoting cell proliferation (Kerber et al., 1998; Sudarsan et al., 2002).

Here we set out to test these alternative models of how the distal MpT is patterned during development, to gain insights into the principles by which secondary axes are generated and patterned.

## Results

### An asymmetric signalling mechanism is unlikely to account for patterning of the distal Malpighian tubules

To assess patterning of the distal MpT, we used markers of the transcription factor Homothorax (Hth), which is expressed in the distal segment of the MpTs (Zohar-Stoopel et al., 2014; Fig. 2A-D), and Dachshund (Dac), which we found to also be expressed in this segment (Fig. 2E-F). These genes are likely to be involved in normal differentiation of the distal cells, leading to their specialised structure and physiological function. Indeed, Hth has been implicated in correct positioning of the kink in each anterior MpT (Zohar-Stoopel et al., 2014), and we found that following depletion of Hth by RNAi, excessive uric acid can be observed in the adult MpT, suggesting a disruption in normal physiological function (Fig. S1).

**Figure 2.**
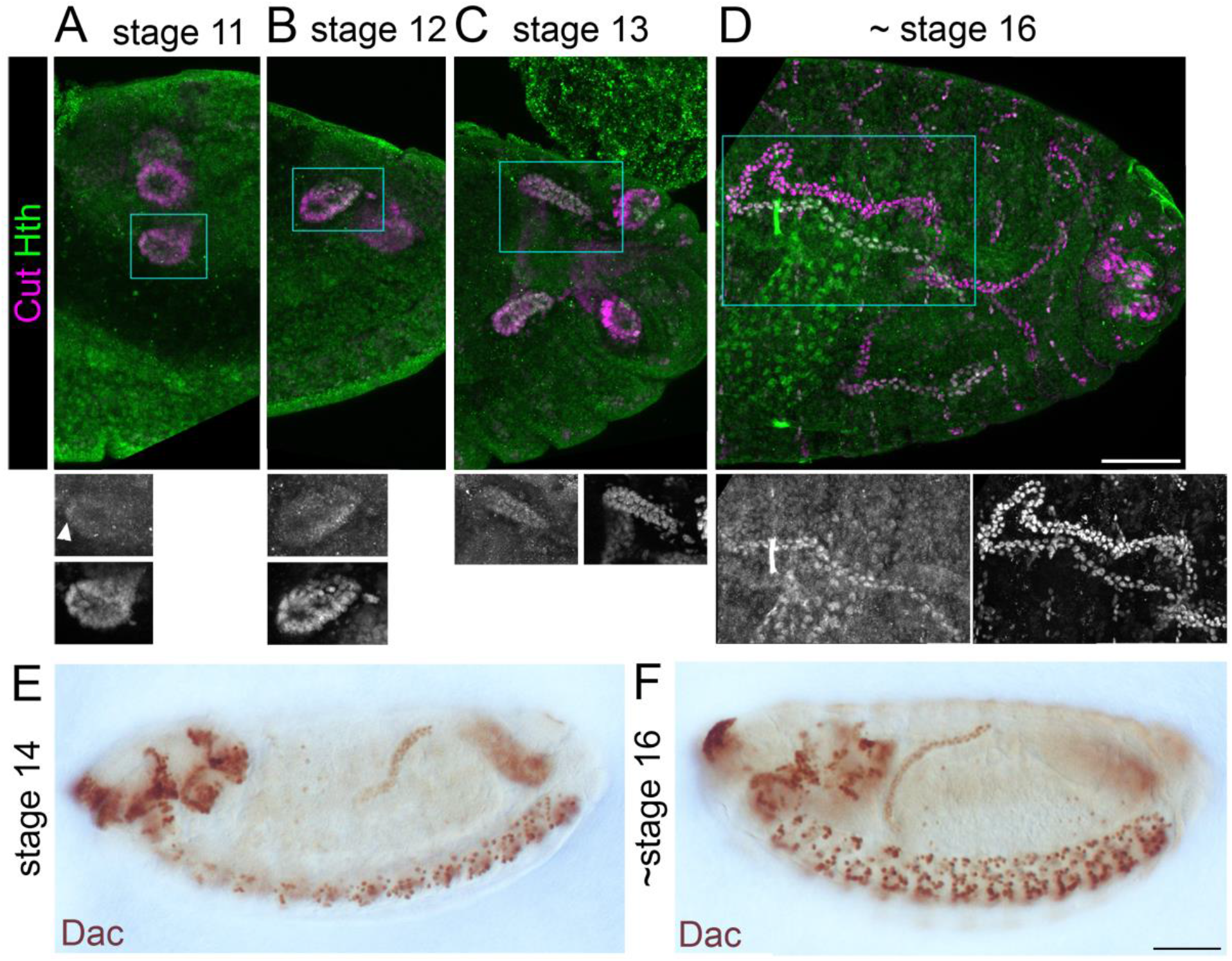
Homothorax and Dachshund are expressed in a distal domain of developing Malpighian tubules. **(A-D)** Stage 11-16 embryos (as indicated) stained with anti-Cut to show MpT nuclei, and with anti-Homothorax (Hth). Homothorax starts to be detectable in nuclei of the distal MpT from about stage 11 (arrowhead in A). Scale bar = 50μm. **(E-F)** Stage 14-16 embryos (as indicated) stained with anti-Dachshund. Scale bar = 50μm.

We then used genetic approaches to manipulate EGF and Wg signalling. We drove expression of constitutively active EGFR (*UAS-λtop*) or dominant negative EGFR (*UAS-EGFR-DN*) in the developing tubules using *CtB-Gal4* which expresses in all MpT cells from the stage when they bud from the gut (SudarsaIn et al., 2002). If EGF signalling is required to induce distal identity it would be expected that the constitutively active form would induce expression of distal markers throughout the MpT, whilst distal marker expression would be abolished by the dominant negative form. Distal tubule identity, indicated by the pattern of Hth expression, was normal in both of these conditions (Fig. 3A-C). This argues against a mechanism whereby EGF signalling is required for distal MpT cell identity.

**Figure 3.**
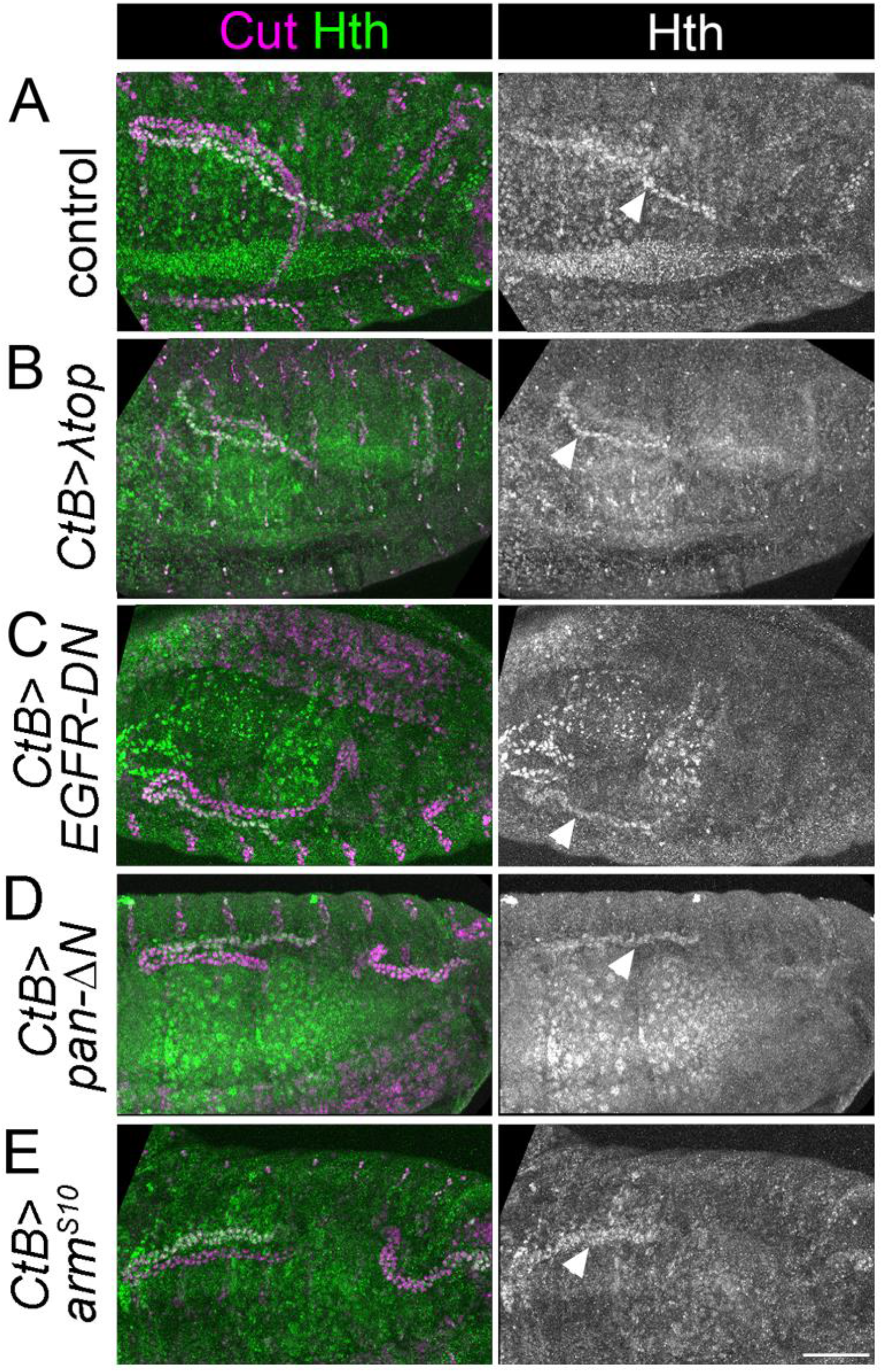
EGF or Wg pathway activity do not appear to pattern the distal Malpighian tubule. **(A-E)** ~Stage 16 embryos stained to show MpT nuclei (anti-Cut) and the nuclei of cells from the distal MpT segment (anti-Hth, arrowheads). In each example at least one of the pair of anterior MpTs is clearly visible. **(A)** Control (*w1118*) embryo. **(B)** *CtB-Gal4>UAS-λtop* embryo. **(C)** *CtB-Gal4>UAS- EGFR-DN* embryo. **(D)** *CtB-Gal4>UAS-pan/dTCFΔN* embryo. **(E)** *CtB>UAS-armS10* embryo. Scale bar = 50μm.

We also tested the effect of driving expression of a construct which inhibits Wg signalling (*UAS-pan/dTCF∆N*) or which constitutively activates Wg signalling (*UAS-arm*^*S10*^) using *CtB-Gal4*. We previously reported that both of these genotypes disrupt the normal proximal patterning of the MpTs (Beaven and Denholm, 2018). If Wg signalling is required to repress distal identity it would be expected that distal identity markers would be expressed throughout the MpT upon inhibition of Wg signalling, and that their expression would be lost when constitutively activating Wg signalling. However we did not see any alteration in the pattern of Hth expression (Fig. 3A,D,E). Therefore we find no evidence that Wg in the proximal MpT acts to inhibit distal tubule identity.

These experiments extinguish EGF and Wg signalling pathways, the two strongest candidate pathways, as signals emanating from one or other end of the MpT to define distal identity. However they do not rule out the possibility that other signalling pathways could be involved in such a mechanism.

### Acquisition of distal tubule identity requires Wingless

In contrast to our findings when manipulating Wg using reagents driven by the *CtB-Gal4* driver, we observed that in a null mutant of *wg* (*wg*^*1-17*^), expression of Dac in the MpTs is abolished (Fig. 4B). Note that MpTs are much shorter in this mutant, as previously reported (Skaer and Martinez Arias, 1992), relating to roles of Wg in specifying the tip-cell and driving cell proliferation. This seeming contradiction to the data reported above could be explained by the timing of Wg manipulation when using the *CtB-Gal4* driver line. The *CtB* enhancer is expressed in the developing MpTs from stage 11 or earlier, and would be expected to be expressed from the very start of their development (Jack and DeLotto, 1995; personal communication with Flybase - Skaer, H. (2002.8.7). CtB-Gal4 line; Sudarsan et al., 2002), however it may be that the Gal4 protein does not reach sufficient levels to induce good inhibition of Wg signalling until later in MpT development. It therefore seems plausible that Wg participates in specifying distal identity early on in MpT development, in the period before *CtB-Gal4* is effective. This notion is also supported by the normal size of the MpTs when manipulating Wg using the *CtB-Gal4* driver, suggesting that Wg’s early roles, for example in cell division, are not significantly perturbed by manipulations using *CtB-Gal4*.

**Figure 4.**
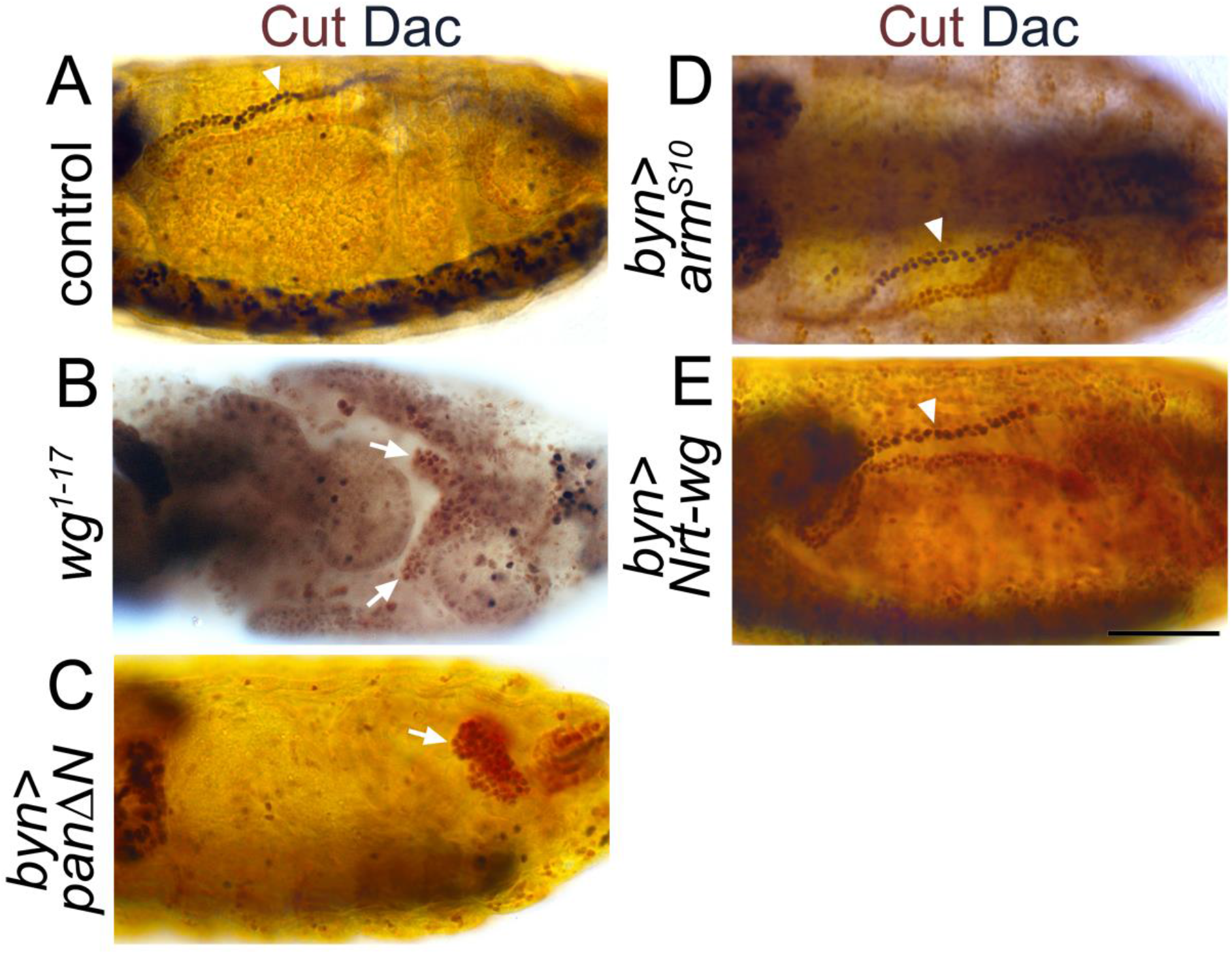
Wingless is required for expression of distal tubule markers. **(A-E)** ~Stage 16 embryos stained to show MpT nuclei (anti-Cut, brown) and the nuclei of cells from the distal MpT segment (anti-Dac, black, arrowheads). Arrows indicate MpTs, in cases where anti-Dac staining is absent. **(A)** Control embryo (*w1118*). **(B)** Homozygous *wg1-17* embryo. **(C)** *byn-Gal4>UAS-pan/dTCFΔN* embryo. **(D)** *byn-Gal4>UAS-armS10* embryo. **(E)** *byn-Gal4>UAS-Nrt-wg* embryo. Scale bar = 50μm.

To further test the hypothesis that Wg acts in early MpT development, we repeated the *UAS-pan/dTCF∆N* experiment using *brachyenteron-Gal4* (*byn-Gal4*), which is expressed in the proctodeum in the shared primordia that gives rise to the hindgut and MpTs, and subsequently in the developing MpTs (Hatton-Ellis et al., 2007; Weavers and Skaer, 2013). This produced a similar result to *wg*^*1-17*^ in that MpTs were greatly reduced in size, and distal identity, as indicated by Dac expression, was abolished (Fig. 4A-C). Wg is therefore required to establish distal identity.

We also tested whether the distal tubule domain could be expanded by ectopic activation of Wg signalling. We expressed either *UAS-arm*^*S10*^ or membrane tethered Wg (*UAS-Nrt-wg*), using *byn-Gal4* (Nrt-Wg is known to function at least as well in activating Wg signalling as untethered wild-type Wg (Beaven and Denholm, 2018; Chaudhary et al., 2019; Zecca et al., 1996)). Neither of these manipulations altered the domain of Dac expression (Fig. 4A,D,E). Together, these data show Wg function is necessary but not sufficient for distal tubule identity. The lack of sufficiency suggests other factors act in concert with Wg to establish distal identity.

Our findings therefore support a model that Wg acts to specify distal cell identity in the MpT primordium during the earliest stage of their development (Fig. 1B). However alternative hypotheses could also explain this phenotype. Firstly, we have not ruled out a signal other than Wg emanating from the gut to repress distal MpT gene expression (Fig. 1 A”). This would be another way to explain the loss of distal identity in the *wg* mutant tubules, as the tubules are much smaller which could bring all the MpT cells within range of the putative inhibitory signal. We previously found that *crossveinless c* mutant tubules (*cv-c*^*c524*^) are morphologically abnormal and do not extend far from the gut (Denholm et al., 2005), providing an ideal means to test this hypothesis. We found that regions of these short *cv-c* mutant tubules expressed the distal markers Hth and Dac (Fig. 5A,B). This provides further evidence against a model for an inhibitory signal emanating from the proximal end of the MpT to inhibit distal identity (Fig. 1A’’), and also indicating that such an inhibitory signal cannot account for the loss of distal identity in the shortened *wg* mutant.

**Figure 5.**
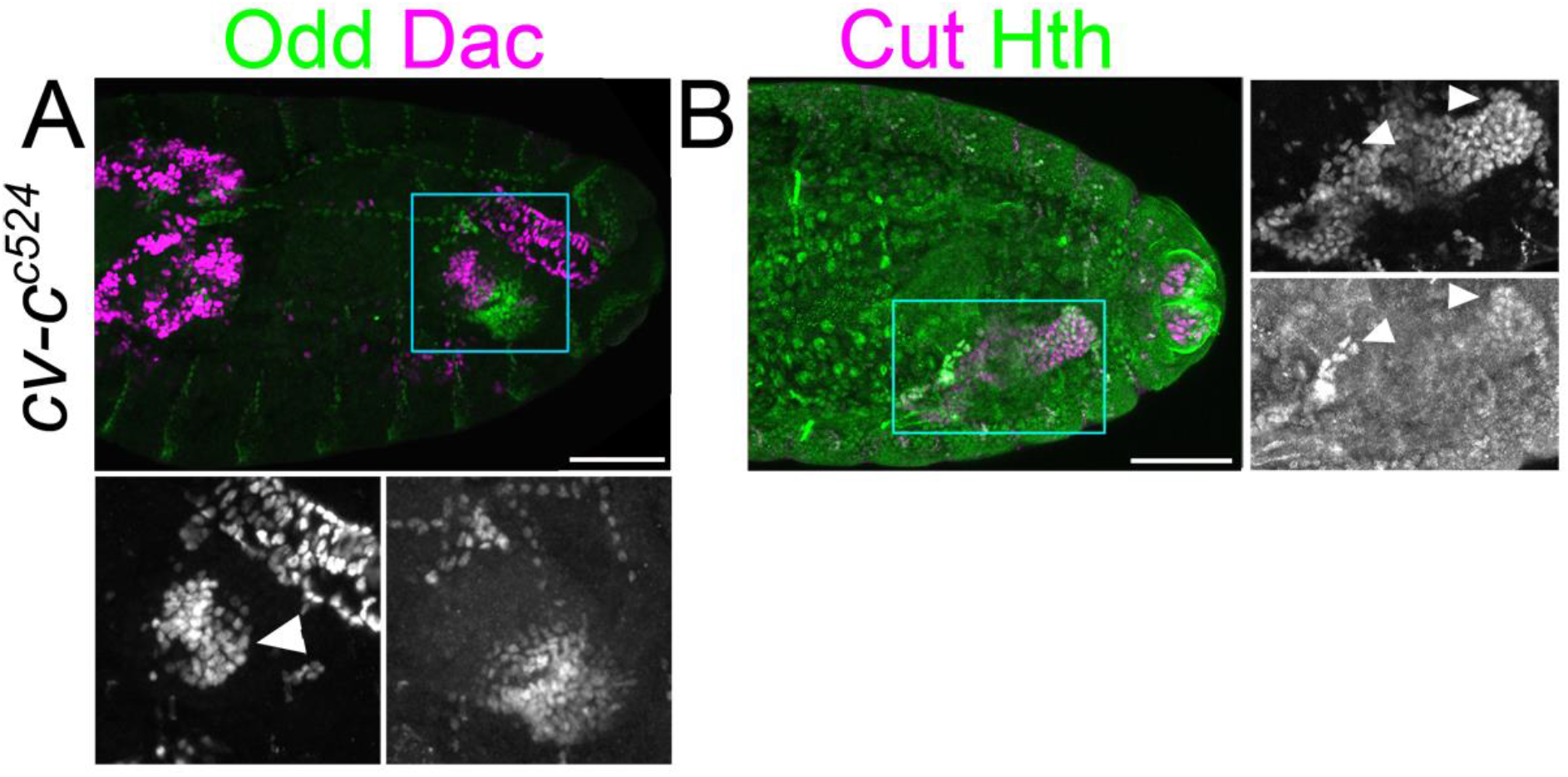
Reducing Malpighian tubule length does not lead to inhibition of distal transcription factor expression. **(A-B)** ~Stage 15 embryos. **(A)** Homozygous *cv-cc524* embryo stained to show the nuclei of the proximal MpT cells (anti-Odd skipped) and nuclei from the distal MpT cells (anti-Dac, arrowhead). **(B)** Homozygous *cv-cc524* embryo stained to show the nuclei of the MpT cells (anti-Cut) and nuclei from the distal MpT cells (anti-Hth, arrowheads). Scale bars = 50μm.

### Cell proliferation is required to expand the distal population of Malpighian tubule cells

A second alternative hypothesis to explain the loss of Dac expression in the *wg* mutant MpTs relates to the role of the tip- and sibling-cells. As well as producing the EGF ligand Spitz (Saxena et al., 2014), the tip-/sibling-cell(s) could act to produce other signals which induce distal MpT identity. It is known that Wg is required for formation of the tubule tip/sibling-cells through its role in activating expression of *achaete-scute complex* (*as-c*) genes (Wan et al., 2000), so it is conceivable that the loss of Dac expression in the *wg* mutant is an indirect consequence of losing the tip and sibling-cells.

We therefore analysed embryos mutant for the *as-c* genes (*Df(1)sc-B57*) which fail to develop the tip/sibling-cells (Hoch et al., 1994; Sudarsan et al., 2002). In mutant embryos, the MpTs still expressed distal transcription factors (Fig. 6A-C). This experiment is informative in two ways: it provides clear evidence against a requirement for the tip-cell lineage in specifying distal identity downstream to Wg, i.e. it points to a more direct role for Wg in specifying distal identity. Secondly, it provides further evidence against the model of distal identity induction by a signal originating from the tip/sibling-cells (Fig. 1A’).

**Figure 6.**
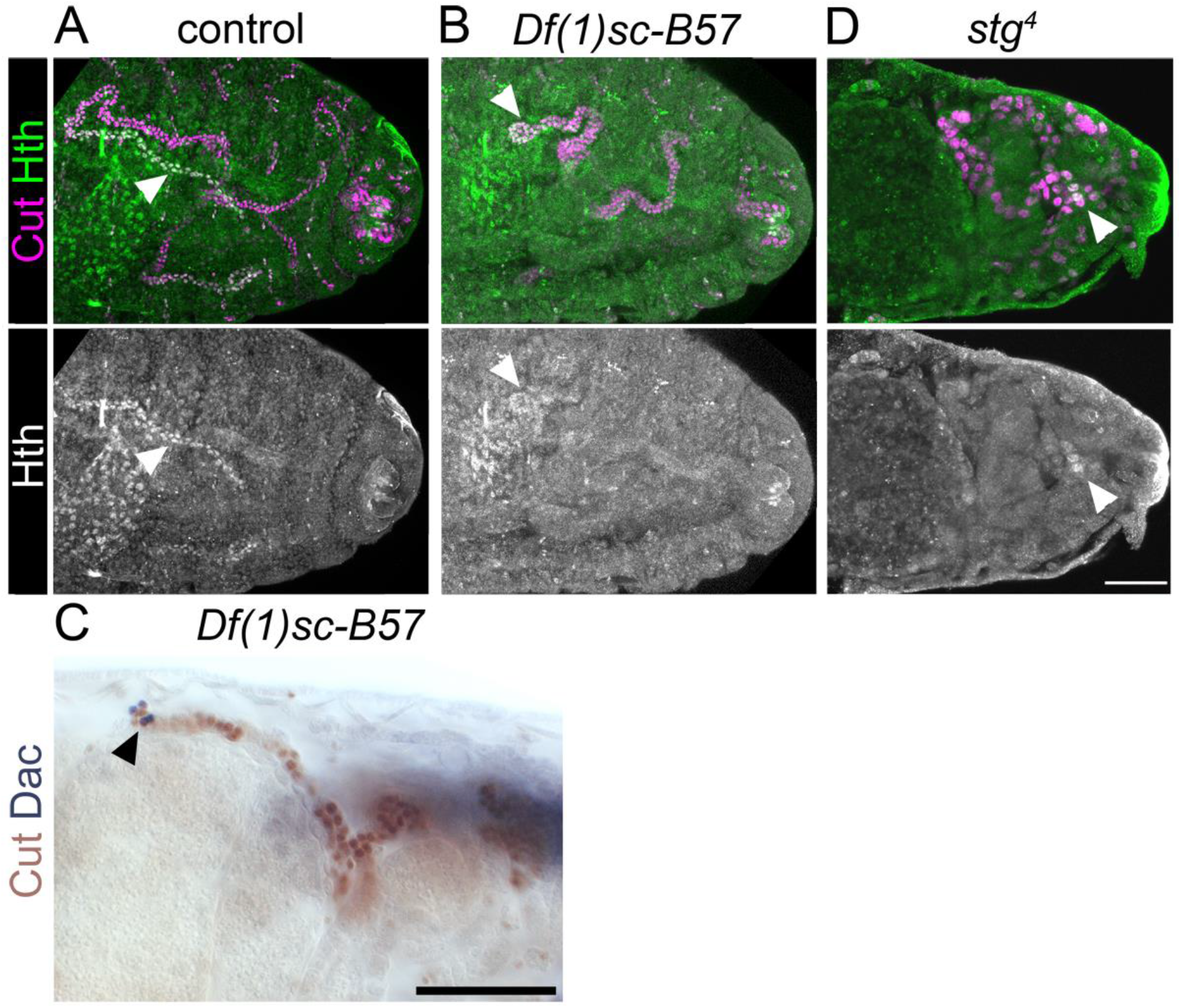
*achaete-scute complex* genes are not required to specify distal tubule identity, but are required to expand the pool of distal identity cells. **(A-B** and **D)** ~Stage 16 embryos stained to show the nuclei of the MpT cells (anti-Cut) and nuclei from the distal MpT cells (anti-Hth, arrowheads). **(A)** Control (*w1118*) embryo. **(B)** Homozygous *Df(1)sc-B57* embryo. **(C)** ~stage16 homozygous *Df(1)sc-B57* embryo stained to show MpT nuclei (anti-Cut, brown) and the nuclei of cells from the distal MpT segment (anti-Dac, black, arrowhead). **(D)** Homozygous *stg4* embryo. Scale bars = 50μm.

Intriguingly, Dac and Hth are expressed only in a small cluster of cells at the distal tubule end, in the *as-c* mutant embryos (Fig. 6A-C). We considered the best explanation of this result to be that the distal identity is specified in a small population of tubule cells early in tubule development, with this pool of cells then being expanded (during stage 12) by cell proliferation driven by the tip-cell. If this hypothesis is correct, then blocking cell division should also reduce the number of tubule cells expressing distal transcription factors. In *string* mutant embryos (*stg*^*4*^) cell division is arrested in cycle 14 (Edgar and O’Farrell, 1990). We found that only a very small number of tubule cells expressed Hth in this mutant (Fig. 6D), which supports our hypothesis of early specification, and subsequent amplification, of distal identity cells.

### Wingless is required early in MpT development to specify distal identity, and appears to act in an autocrine and/or juxtacrine manner

As a direct test of the hypothesis that Wg is required in the tubule primordium in the very early stages of tubule development, we made use of a temperature sensitive mutant (*wg*^*1-12*^), which has previously been used to map the temporal requirements of Wg in tubule development (Beaven and Denholm, 2018; Skaer and Martinez Arias, 1992). In batches of *wg*^*1-12*^ embryos which had developed at the restrictive temperature for 230-290 minutes, before switching to the permissive temperature for the rest of development, almost all tubules lacked Dac expression (Fig. 7A). In batches of *wg*^*1-12*^ embryos which had developed at the restrictive temperature for 210-250 minutes before shifting to the permissive temperature, approximately half of the embryos had tubules lacking Dac expression whilst in the other half Dac expression appeared normal (Fig. 7B). Therefore shifting to the restrictive temperature at ~230 minutes or later abolishes Dac expression. Taking into account that Wg function is considered to take ~25 minutes to recover after shifting to a permissive temperature (Skaer and Martinez Arias, 1992), this suggests that Wg is required until ~255 minutes into embryogenesis to confer distal tubule identity, and that after this time Wg is dispensable. This equates to late stage 9 or early stage 10. In line with our pre-patterning hypothesis (Fig. 1B), we have found evidence that the requirement of Wg in conferring distal tubule identity is during the earliest point of MpT development: in the tubule primordium before the tubules bud out from the gut. The time of Wg requirement is also comparable to its requirement for allocation of the tip/sibling-cells, early in the development of the MpT (Wan et al., 2000).

**Figure 7.**
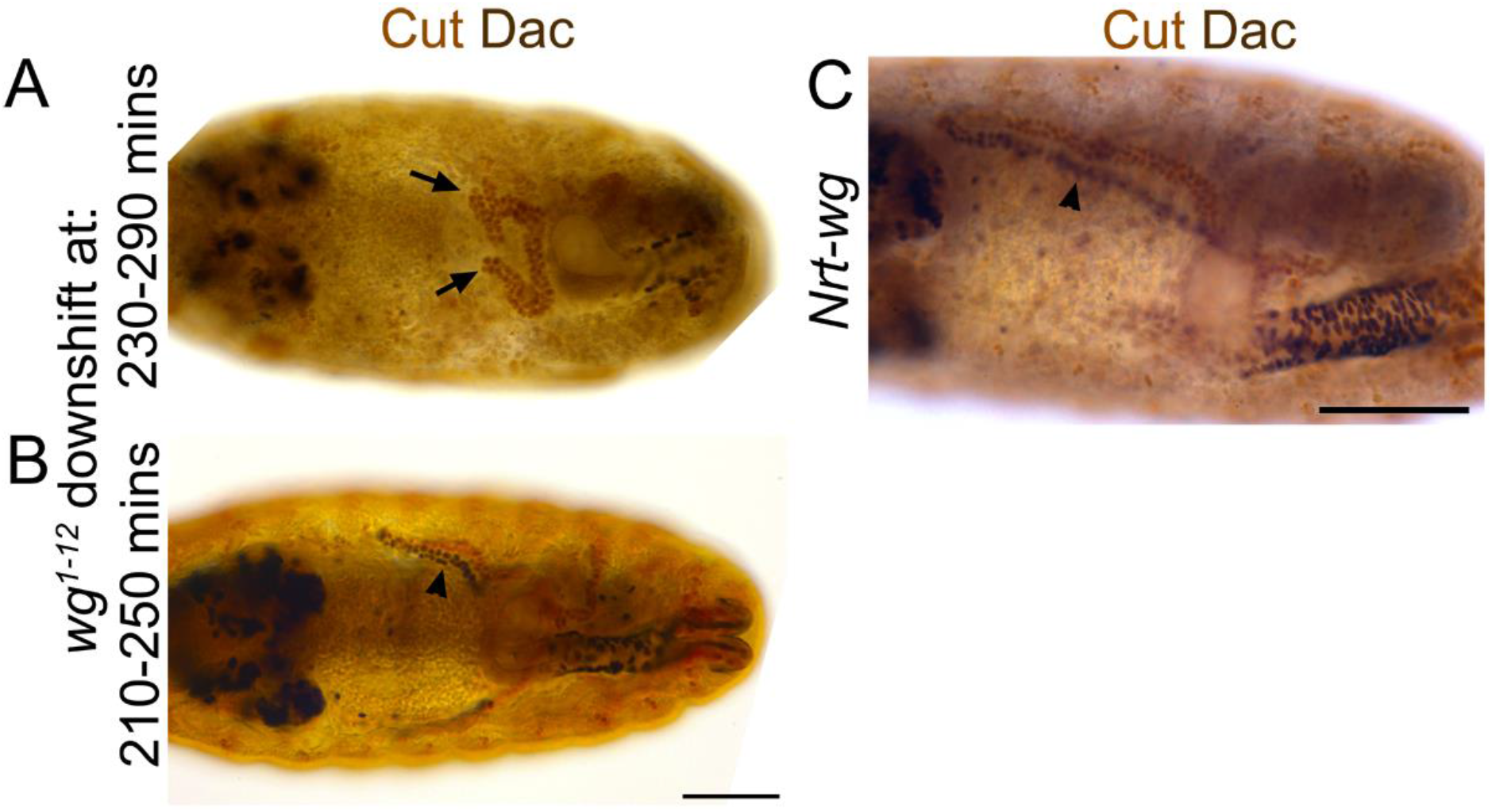
Wingless is only required during early tubule formation for specification of distal tubule identity. **(A-C)** ~Stage 15-16 embryos stained to show MpT nuclei (anti-Cut, brown) and the nuclei of cells from the distal MpT segment (anti-Dac, black, arrowheads). **(A)** Homozygous temperature sensitive allele of *wg* (*wg1-12*) raised at the restrictive temperature until 230-290 minutes when shifted to the permissive temperature. Arrows indicate MpTs which are lacking expression of Dac. **(B)** Homozygous temperature sensitive allele of *wg* (*wg1-12*) raised at the restrictive temperature until 210-250 minutes when shifted to the permissive temperature. **(C)** Homozygous embryo in which endogenous *wg* has been replaced with *Nrt-wg*. Scale bar = 50μm.

We were curious to know whether the role of Wg in specifying distal identity in the MpT primordium requires release and spread of the Wg ligand, as we had demonstrated a requirement for Wg dispersal in patterning the proximal tubule (Beaven and Denholm, 2018). Therefore we assessed the expression pattern of Dac in tubules using a fly line in which endogenous Wg is replaced with a non-diffusible membrane tethered form (*Nrt-wg*) (Alexandre et al., 2014). We found that the pattern of Dac expression appears normal in *Nrt-wg* tubules (Fig. 7C). This suggests that the requirement for Wg in establishing distal identity occurs within the cells which express Wg (autocrine signalling) and/or their immediate neighbours (juxtacrine signalling). This contrasts to the role of Wg in patterning the proximal MpT, where release and spread is essential.

### Wingless is expressed in the half of the Malpighian tubule primordium which neighbours the developing hindgut

In order to better understand the role of Wg during MpT development, we made use of a line in which Wg has been tagged with GFP at the endogenous genetic locus (Port et al., 2014). In fixed samples, anti-GFP staining of these embryos gave much clearer staining relative to background than we have seen using anti-Wg (Beaven and Denholm, 2018). We combined this with the tubule marker anti-Cut, and assessed localisation during early tubule development.

At stage 10, we found that Wg-GFP is expressed in the region of the tubule primordium abutting the developing hindgut primordium, but not the region abutting the developing midgut (Fig. 8A). This is consistent with the pattern observed with anti-Wg staining (Wan et al., 2000). This region of the tubule primordium is the region from which the tip mother cell segregates (Wan et al., 2000), and is therefore likely to be the population of cells from which the distal cell lineage is derived. This patch of early Wg expression coincides with the period Wg activity is required to establish distal identity in the tubule (previous section), strengthening the evidence that Wg is the early patterning signal.

**Figure 8.**
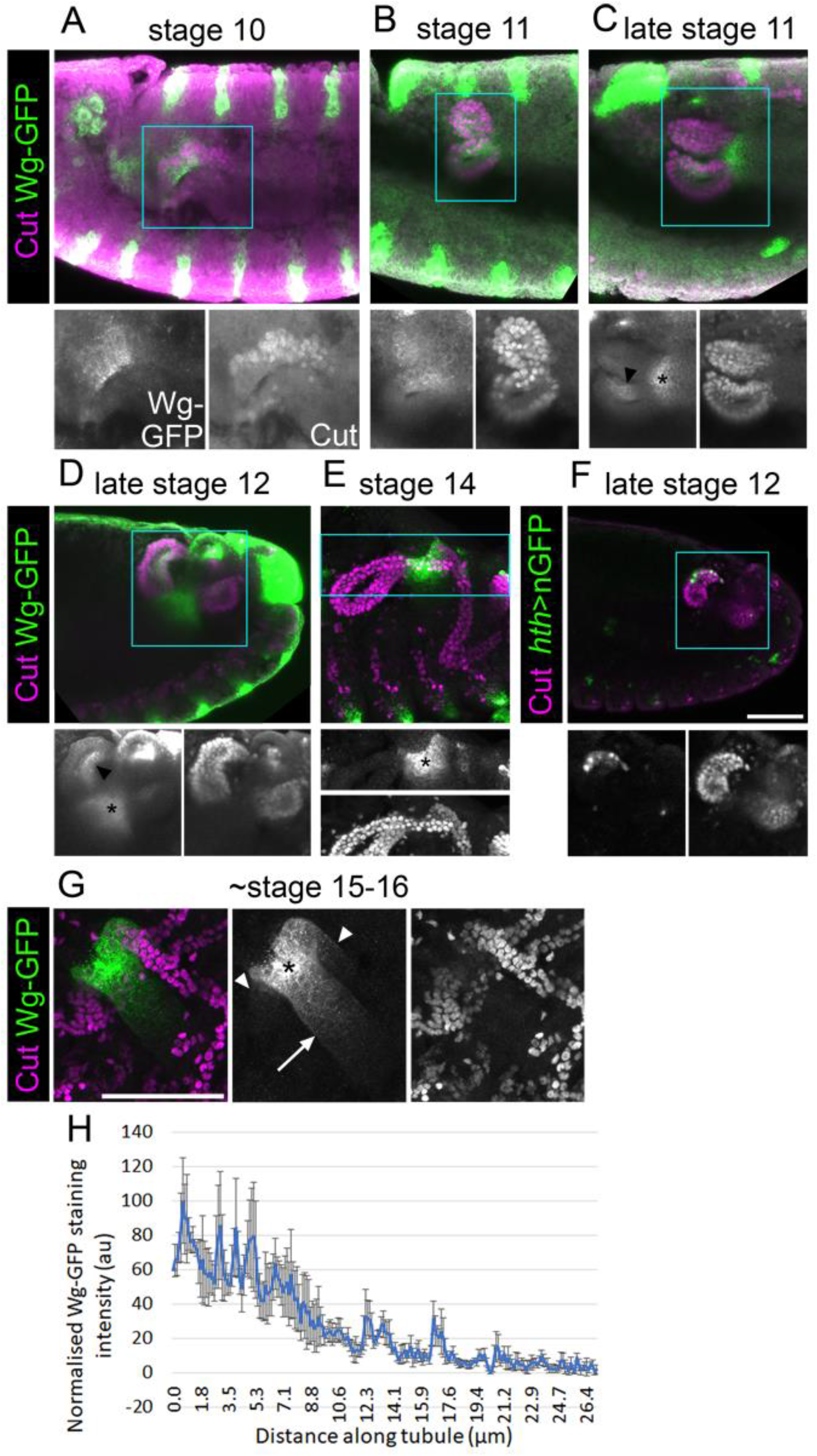
Wingless is present in the half of the tubule primordium that abuts the developing hindgut. **(A**-**E)** Stage 10-14 embryos expressing Wg-GFP, stained with anti-GFP, along with anti-Cut to show nuclei of MpT cells. Wg-GFP can be observed in **(A)** MpT cells closest to the hindgut as the MpTs begin budding out at stage 10, **(B-D)** on one side of the MpTs at stage 11-12 (arrowheads in C + D), **(C-E)** and a gut region near the midgut-hindgut boundary at the proximal MpT ends from late stage 11 (*). **(F)** Labelling of distal MpT cells in late stage 12 *hth-Gal4>UAS-nGFP* embryo stained with anti-GFP, along with staining of MpT cell nuclei (anti-Cut). Scale bar = 50μm (scale bar in F applies to A-F). **(G)** Late stage (~stage 15-16) embryo, expressing Wg-GFP, stained with anti-GFP and anti-Cut. Wg-GFP localises most strongly to a region near the midgut-hindgut boundary (*) from where a gradient extends into the midgut (arrow) and into the two tubule ureters (arrowheads), which branch into the anterior and posterior pairs of MpTs. Scale bar = 50μm. **(H)** A graph of the staining intensity of anti-GFP to show the distribution of Wg-GFP from the proximal tubule end. n=4. Error bars show S.E.M.

## Discussion

We have found evidence of a mechanism in which epithelial tubules, the Malpighian tubules of *Drosophila*, achieve patterning of their P-D axis by early specification of a subgroup of cells which expand during development, giving rise to the distal domain. Wg signalling during early MpT development plays a critical role in specifying their distal identity. This mechanism of early patterning followed by expansion does not rely on dispersal of the Wg ligand. This is in contrast to our previous findings for the role of Wg in patterning the proximal tubule, which does require release and spread of Wg.

In the context of proximal patterning, Wg activates Odd expression to a distance of ~30μm (Beaven and Denholm, 2018). Using Wg-GFP, we find that in late stage embryos (~stage 15-16) Wg in the proximal tubule clearly forms a gradient, which extends ~25μm into the MpT (Fig. 8G,H). It also appears to extend ~40μm into the midgut (Fig. 8G,H). This fits well with the range of Wg dispersal in the wing imaginal disc, which is in the order of ~30μm (Chaudhary et al., 2019), as well as the range of the Wg-GFP gradient observed in the adult gut (Tian et al., 2019). These figures may constitute the upper limit of *Drosophila* Wg dispersal. Considering that the anterior tubules extend to over 300μm by late embryogenesis, and that the domains of distal transcription factors such as Dac and Hth extend to about half this length, this would make the final size of distal expression domains much greater than the effective range of Wg. It seems that the maximum range of Wg is too small to allow Wg scaling to pattern the distal MpT in the manner of a classical morphogen (as, for example, Decapentaplegic (Dpp) does in the wing disc, scaling to 100μm as the disc reaches full size (Teleman and Cohen, 2000; Wartlick et al., 2011)). This could explain why Wg defines the distal tubule domain in the tubule primordium whilst it is a small cluster of cells, and these cells then maintain their distal identity as they proliferate and the tubule elongates.

Our findings that Wg acts in the very earliest stages of MpT development to define distal cell identity also provoked us to contemplate whether the Wg signal is the earliest symmetry breaking step by which the axis is established. Intuitively one might expect the distal cells to arise from the centre of the primordium, as these would naturally become distal as the tubule telescopes out during development. Such a mechanism is responsible for P-D patterning of the *Drosophila* legs; here the discs are patterned as concentric rings, with the centre of the disc ultimately becoming the distal end of the leg (Estella et al., 2012). This patterning is achieved by first specifying the central point of the disc which becomes the distal pole of the P-D axis. This is achieved by Dpp and Wg which are expressed dorsally and ventrally at the anterior-posterior compartment boundary, under regulation of hedgehog from the posterior compartment. The intersection of the Dpp and Wg signals activates Distal-less expression at the centre of the leg disc (Basler and Struhl, 1994; Campbell et al., 1993; Diaz-Benjumea et al., 1994). However what we observe in the MpTs appears different, in that the cells that ultimately give rise to the distal MpT originate on one side of the MpT primordium – the side which abuts the hindgut and expresses Wg (Figs. 1B, 8A), rather than the centre of the primordium. It is conceivable that the established axis of the developing gut acts to break symmetry in the MpT primordium and thus establish the P-D axis of the tubules. In this context the expression of Wg on one side of the MpT primordium is likely to be regulated by the processes concomitantly patterning the developing gut, such as the division of the hind- and midgut. Further processes define a proneural cluster of cells in the developing MpT, and specify the tip-cell lineage within this cluster. These processes are abolished by depletion of Wg in early MpT development (Wan et al., 2000), which may be because only the distal lineage of MpT cells are competent to give rise to the proneural cluster. The emergence of the tip/sibling-cells as a signalling centre could then orchestrate the morphological changes underlying tubule elongation, in a way which results in their surrounding cells coming to have a distal position in the final tubule. The EGF signal which emanates from the tip/sibling-cells drives cell proliferation (Baumann and Skaer, 1993; Kerber et al., 1998; Sudarsan et al., 2002), as well as defining planar cell polarity in surrounding cells to orchestrate their intercalation and the convergent extension of the distal MpT (Saxena et al., 2014), so orchestrating the morphological changes which propel the tip/sibling-cells, and their neighbouring cells with distal identity, to finally reside at the distal end of the extended MpT. Together these processes could convert an already established asymmetry in the gut, into a P-D axis in the MpTs which grow out perpendicular to the gut axis.

The formation of this secondary axis is conceptually similar to the formation of the body’s main axes, in that both are underpinned by the breaking of symmetry under influence from external source(s). It also shows parallels to formation of other studied secondary axes such as *Drosophila* legs, where establishment of the P-D axis requires an initial breaking of symmetry which is derived from patterning of the main body axes. In diverse examples, including all the *Drosophila* imaginal discs, as well as vertebrate limbs, primordia develop straddling compartment boundaries formed along the primary body axes. As the cells of the primordia straddle boundaries, they are conferred a developmental asymmetry from the outset, allowing them to establish secondary axes (Basler and Struhl, 1994; Meinhardt, 1983a; Meinhardt, 1983b; Tabata et al., 1995). This may explain why the MpT primordia arise precisely at the boundary between the developing hindgut and midgut.

The mechanism we have uncovered for patterning the distal tubules, i.e. early patterning and expansion of a distal domain, must require some form of cellular memory. Distal cells appear to begin expressing transcription factors such as Dac and Hth some time after their exposure to the Wg signal, and their expression is maintained into the adult, in the absence of Wg. Therefore, there must be a mechanism for cells to retain their identity in the absence of the initial signal. One means to achieve such cellular memory is transcriptional memory as has been characterised for Hox genes, for example in conferring positional identity along the anterior-posterior axis, as well as for hedgehog expression in the *Drosophila* wing discs. Transcriptional memory of Hox genes and hedgehog involves binding of Trithorax- and Polycomb-group proteins to Trithorax response elements (TREs) and Polycomb response elements (PREs) which are regulatory chromosomal regions (Francis and Kingston, 2001; Maurange and Paro, 2002; Pérez et al., 2011; Wang et al., 2009).

In the case of patterning mechanisms by Wg, it has been shown in wing discs that the frizzled 2 receptor can function to maintain Wg signalling in cells which are no longer exposed to the Wg ligand (Chaudhary et al., 2019). We do not think that this mechanism is the one used in the MpTs, as it relies on perpetuating activity of the Wg signalling pathway (which we found not to be necessary in our experiment driving *dTCF∆N*, Fig. 3D). It has also been shown in gastruloids that cells retain memory of their exposure to Wnt via an unknown mechanism, which controls their responsiveness to activin (Yoney et al., 2018).

To further understand the mechanisms by which Wg could regulate distal target genes, we looked at the activity of *dac* enhancers containing dTCF/Pan binding sites, but found that these enhancers, which regulate expression of *dac* in the leg disc, are not responsible for *dac* expression in the distal tubule (Fig. S2). This provides evidence to suggest regulation by Wg is indirect (at least for *dac*), indicating that Wg might trigger a signalling cascade which activates and maintains expression of distal identity genes. Testing the regulation of Hth may also shed light on this mechanism. A genomic fragment of *hth* was previously found which contains the enhancer region responsible for its expression in the distal MpT (Zohar-Stoopel et al., 2014). There is a dTCF/Pan binding site within this region (Junion et al., 2012), which provides a putative site for direct Wg regulation of *hth* expression in the MpT.

We have found that Wg, acting at different times during development, plays roles in patterning both the distal and proximal ends of the MpT, and is required for the expression of different target genes in these two contexts. This highlights how context dependent the outcomes of Wg signalling are, even within a single tissue. We had previously found that constitutively activating Wg signalling throughout the MpT leads to an expansion of the proximal Odd expression domain, but that the majority of the MpT including all of the distal region still lacked Odd expression (Beaven and Denholm, 2018). In contrast, we report here that activating Wg signalling throughout the MpT from the very beginning of its development does not cause expansion of the distal domain, as indicated by Dac expression (Fig. 4D,E). The different contexts may be underpinned by differences in the signals which proximal and distal MpT cells have encountered during their development, apart from Wg.

Following the early period of expression of Wg in the half of the tubule primordium which abuts the hindgut at stage 10 (Fig. 8A), there appears to be two subsequent and distinct patterns of Wg expression. Firstly, during stage 11-12 Wg is expressed on one side of the MpT (Fig. 8B-D). This is not the distal cell population, which becomes clear when comparing to expression of distal markers at stage 12 (Fig. 8F). This expression on the inner side of the MpT is consistent with a previous report of *wg* mRNA and Wg protein expression at this stage, which is considered to be driving cell division in this cell population (Skaer and Martinez Arias, 1992). It therefore seems likely that this represents a second, functionally distinct phase of Wg expression. Secondly, from late stage 11, throughout the rest of embryogenesis, we observed a third distinct region of Wg expression in the developing gut abutting the proximal tubule (Fig. 8C-E and G). This is the region of Wg that specifies the identity of a proximal tubule domain, being required for expression of Odd skipped (Odd) (Beaven and Denholm, 2018). This expression of Wg at the midgut-hindgut boundary persists stably through larval and adult stages, where it regulates cellular behaviours of the gut and MpT (Fang et al., 2016; Takashima et al., 2008; Tian et al., 2016; Tian et al., 2019; Xu et al., 2018).

Context dependence would allow Wg to play a complex and rapidly changing set of roles in MpT development, in specifying distal identity in the primordium, driving cell proliferation on one side of the tubule and in specifying proximal identity in the later MpT. These multiple, sequential phases of Wg activity are reminiscent of findings from the wing discs. Here the first phase of Wg function is to subdivide the identity of the disc, by contributing to specification of wing fate as opposed to wing hinge and notum fate. This occurs during the larval second instar, during which Wg is expressed in a wedge of cells in the ventral/anterior compartment (Couso et al., 1993; Klein and Arias, 1998; Ng et al., 1996; Williams et al., 1993). Wg is expressed in the entire wing primordium until early third instar larvae where it contributes to growth and patterning of the disc, but is not expressed in the rest of the wing disc. By the late third larval instar Wg is only expressed in a narrow line of cells at the presumptive wing margin, in a double ring around the distal region of the wing and in part of the notal region (AlexandIre et al., 2014; Couso et al., 1993; Couso et al., 1994; Martinez Arias, 2003; Ng et al., 1996; Williams et al., 1993). The roles of Wg at the presumptive wing margin include patterning the wing (Chaudhary et al., 2019; Swarup and Verheyen, 2012 -references within).

The findings here provide mechanistic insights into how a secondary axis is established and patterned in an epithelial tubule. Epithelial tubules are a fundamental component of many organs, and are frequently patterned along their P-D axis. The oviduct or fallopian tube is one example of this, and recent evidence shows that distinct distal and proximal populations of oviduct cells are specified early in development before the oviduct has extended by cell proliferation. These two cell populations form distinct stable lineages (Ford et al., 2021). It may therefore be that early patterning followed by tissue expansion could be a widespread means to pattern the P-D axis of epithelial tubules. Such a mechanism may be important in patterning the P-D axis of nephrons in the kidney, which are also known to require Wnt signalling (Deacon et al., 2019; Lindstrom et al., 2015).

## Materials and Methods

### Fly stocks

Flies were cultured on standard media at 25°C unless otherwise stated. Control flies were from a *w*^*1118*^ or Oregon-R stocks. Mutations were balanced over YFP bearing chromosomes (*dfId\ -GMR-nvYFP* balancers (Le et al., 2006)), in order that homozygous mutant embryos could be selected. The following stocks were also used: *wg-GFP* (Port et al., 2014), *CtB-Gal4;CtB-Gal4/TM3* (Sudarsan et al., 2002), *byn-Gal4* (gift from R. Reuter), *UAS-λtop* (gift from T. Schüpbach), *UAS-EGFR-DN* (#5364, BSC), *UAS-pangolin/dTCF∆N* (van de Wetering et al., 1997), *UAS-armadillo*^*S10*^ (Pai et al., 1997), *wg*^*1-17*^ (Baker, 1987), *cv-c*^*C524*^ (Denholm et al., 2005), *Df(1)sc-B57* (González et al., 1989), *stg*^*4*^ (#2500, BSC), *hth-Gal4* (212052/VT, VDRC), *UAS-nGFP* (Shiga et al., 1996), *dac RE-lacZ* (Giorgianni and Mann, 2011), *∆PRE2* (Ogiyama et al., 2018), *byn*^*5*^ (Singer et al., 1996), *srp*^*3*^ (Jürgens et al., 1984), *UAS-hth-dsRNA* (TRiP line HMS01112), *Nrt-wg*, a *wg* mutant bearing a transgenic insertion of *wg* fused to *Nrt* (*wg[KO; NRT–Wg; pax-Cherry]* (Alexandre et al., 2014)), *UAS-Nrt-wg* (gift from J-P. Vincent), and a temperature sensitive allele of *wg* (*wg*^*1-12*^ (Nüsslein-Volhard et al., 1984)). For temperature shift experiments with *wg*^*1-12*^ a restrictive temperature of 25°C was used for depletion of Wg and a permissive temperature of 18°C was used for recovery of Wg levels, with timings as indicated in the text and Fig. 7A-B.

### Embryo fixation and staining

Embryos were collected on grape juice agar plates with yeast paste. Embryos were fixed and antibody stained using standard techniques (Weavers and Skaer, 2013). Antibodies were diluted in, and washing steps performed with, PBST + BSA (PBS with 0.3% Triton X-100 and 0.5% bovine serum albumin). The following antibodies were used: anti-Cut (mouse, 1:200, 2B10-c from DSHB), anti-GFP (goat, 1:500, ab6673, from Abcam), anti-Futsch (22C10 from DSHB) which marks the tip-cell (Hoch et al., 1994), either of two anti-Hth antibodies which produced comparable results (rabbit, 1:1000, AS1924, (Kurant et al., 1998)) or (goat, 1:100, (dG-20) sc-26187 from Santa Cruz), anti-Dac (mouse, 1:100, mAbdac1-1-c from DSHB), anti-β-galactosidase (rabbit, 1:10000, ICN Biomedicals), anti-Odd skipped (rabbit, 1:400, gift from J. Skeath). For fluorescence stainings, secondary antibodies from Jackson ImmunoResearch of the appropriate species tagged with 488, Cy3 or Cy5 fluorophores were used, and embryos were mounted in Vectashield (Vector Laboratories) or 85% glycerol, 2.5% propyl gallate. In cases where DAB staining was used, following incubation with the first primary antibody, embryos were incubated for 1hr with a biotinylated antibody of the appropriate species, washed 3x 10 minutes, incubated for 30 mins in ABC solution (Vector Elite ABC kit, Vector Laboratories), washed 3x 10 minutes and then incubated in 600μl PBST to which was added 30μl of 10mg/ml DAB, 30μl 0.06% H202, and 60μl of 1% Nickel(II) chloride in order to develop a black stain. After rinsing 3x in PBST the second primary antibody was added, and the DAB staining protocol outlined above was repeated, except that Nickel(II) chloride was not added so that a brown stain developed. After rinsing 3x in PBST, embryos were dehydrated with 5 minute incubations in 50%, 70%, 90% and 3x 100% ethanol. Embryos were left overnight in Histoclear, rinsed 2x in acetone and mounted in DPX new (Merck).

### Adult MpT analysis

Tubules were dissected from young (<10 day old) adults, and viewed immediately (without fixation) on a slide in PBS.

### Imaging and image analysis

Fluorescence images were taken using either a Nikon A1R or Zeiss LSM800 confocal microscope. Maximum intensity projection images were generated using Fiji. Images of DAB-stained embryos were imaged with a Zeiss Axioplan light microscope with a Leica DFC 425C camera. Fiji was used for quantification of Wg-GFP localisation along the MpT. Grey values were taken from maximum intensity projections of the anti-GFP images from ~stage 15-16 embryos, along a line from the gut at the proximal tubule end, extending along a MpT. Following background subtraction, the intensity values were normalised to give the brightest pixel a value of 100. Means were calculated for each pixel position and finally the data was adjusted to give a maximum staining intensity of 100 (arbitrary units).

## Acknowledgements

We are very thankful to the following people and organisations for their help. For fly stocks: Richard Mann and Anthony Gilmore (*dac RE-lacZ*), Giacomo Cavalli and Bernd Schuttengruber (*∆PRE2*), Fillip Port (*wg-GFP*), Jean-Paul Vincent and Joachim Kurth (*Nrt-wg*), Rolf Reuter (*byn-Gal4*), Trudi Schüpbach (*UAS-λtop*), the Bloomington Drosophila Stock Center and Resource Center (National Institutes of Health grants P40OD018537 and 2P40OD010949-10A1) and the Vienna Drosophila Resource Center (VDRC). For the Odd skipped antibody, James Skeath. For assistance with imaging, Anisha Kubasik-Thayil and Marta Czapranska from the IMPACT imaging facility, University of Edinburgh. For useful advice and reagents, Kyra Campbell, members of Andrew Jarman’s lab and the Edinburgh fly club.

---

**Figure S1.**
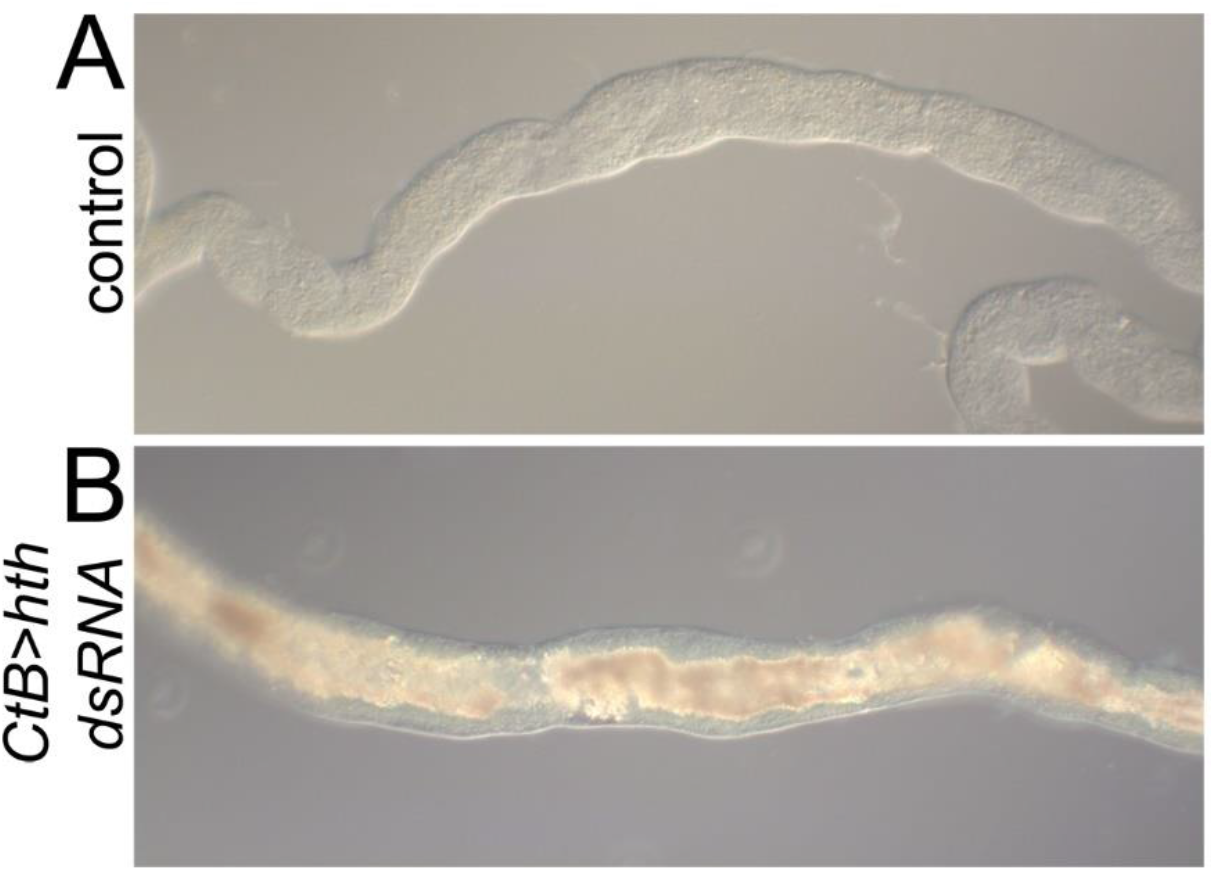
Excessive uric acid accumulates in adult Malpighian tubules following *homothorax* knockdown. **(A)** Section of a control (OrR) adult MpT. **(B)** Section of a *hth* RNAi (*CtB>hth dsRNA*) adult MpT.

**Figure S2.**
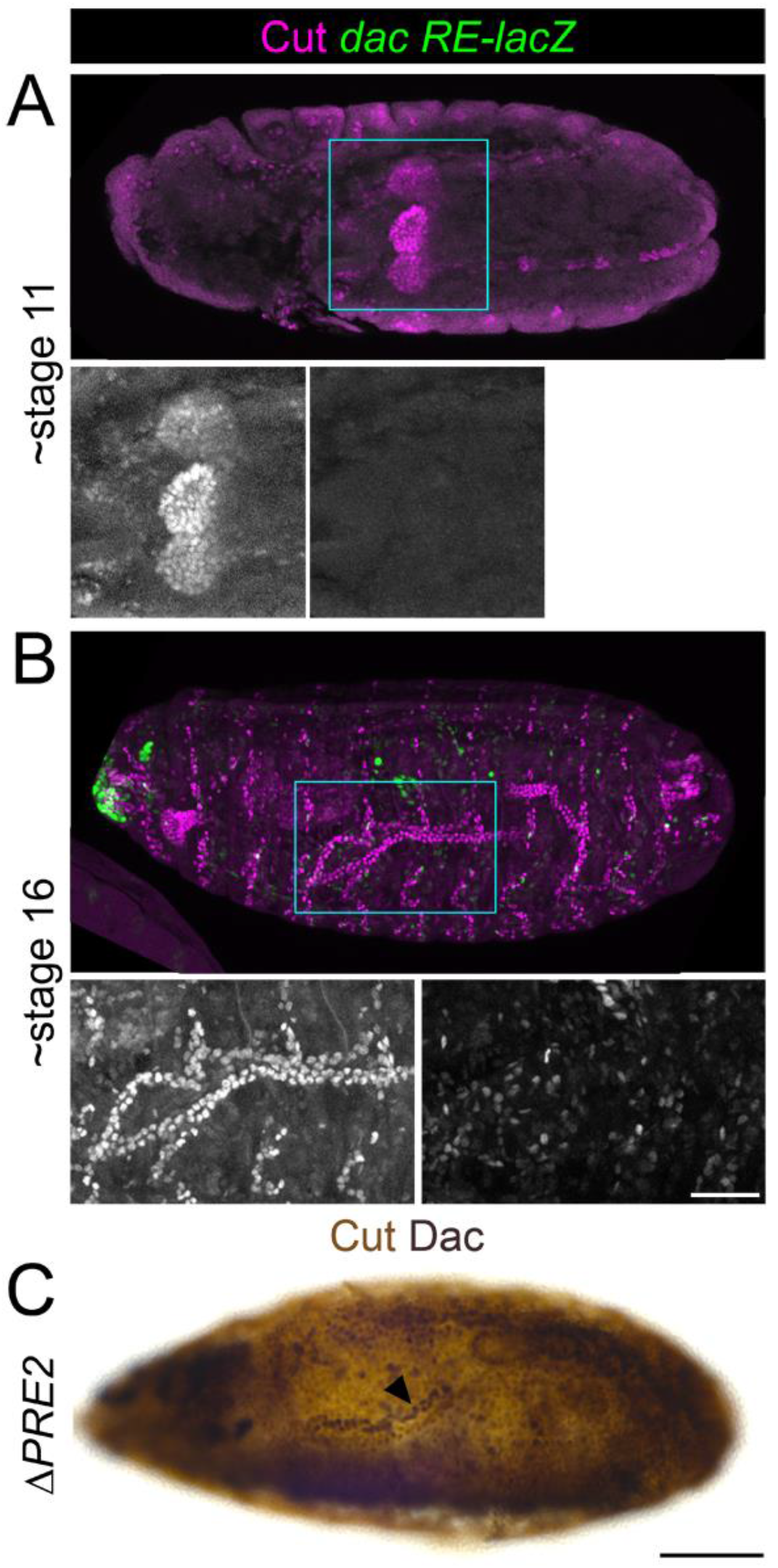
*dac RE* and *dac PRE2* enhancers do not regulate Dachshund expression in the developing Malpighian tubules. **(A-B)** *dac RE-lacZ* embryos stained with anti-β-galactosidase, along with anti-Cut to mark the MpT nuclei. This *dac* enhancer is characterised as driving expression in the leg disc, and contains putative dTCF/Pan binding sites (Giorgianni and Mann, 2011). **(A)** ~stage 11. **(B)** ~stage 16. (**C**) ~Stage 16 homozygous mutant embryo lacking the *PRE2* regulatory region of*dac* (*ΔPRE2*) stained to show MpT nuclei (anti-Cut) and the nuclei of cells from the distal MpT segment (anti-Dac, arrowhead). *PRE2* is a Polycomb response element which has been characterised as a regulatory region for *dac* during leg development (Ogiyama et al., 2018). We were interested in this site as ChIP-on-chip from *Drosophila* embryos showed it to contain a dTCF/Pan binding site (Junion et al., 2012). Scale bars = 50μm.

